# Modelling, Analysis, and Optimization of Three-Dimensional Restricted Visual Field Metric-Free Swarms

**DOI:** 10.1101/2021.05.24.445404

**Authors:** Qing Li, Lingwei Zhang, Yongnan Jia, Tianzhao Lu

## Abstract

Models of collective behaviour have been proved helpful in revealing what mechanism may underlie characteristics of a flock of birds, a school of fish, and a swarm of herds. Recently, the metric-free model gradually occupies a dominant position in the research field of collective intelligence. Most of these models endow every single individual with the ability of a global visual field, which can offer each particle sufficient external information. In this paper, we mainly focus on whether the global visual field is necessary to form a consistent and cohesive group or not. Inspired by the biological characteristic of starlings, we develop a three-dimensional restricted visual field metric-free(RVFMF) model based on Pearce and Turner’s previous work. We further investigate several vital factors governing the convergent consistency of the RVFMF model with the assistance of extensive numerical simulations. According to the simulation results, we conclude that the best view angle of each particle in a swarm increases with the expansion of the population size. Besides, the best view angle gradually becomes stable around 155 degrees when the population size is larger than 1000. We also offer quantitative analysis data to prove that a flock of birds could obtain better consistency under optimal restricted visual field than under global visual field.

## 1. Introduction

Collective intelligence offers an effective solution for groups of organisms during migration, foraging, or predator avoidance, especially for these individuals who cannot survive alone. Scientists are unlocking the secrets behind animals’ collective behaviour using high-speed video and software to figure out how they do it. Generally speaking, models of collective behaviour have proved helpful in revealing what mechanism may underlie characteristics of a flock of birds, a school of fish, and a swarm of herds.

In 1986, Reynolds firstly introduced the Boid model to depict the emersion of actual collective behaviour of birds in a computer animation simulation, which has been regarded as the starting point of scientific research on the collective model [1]. Since then, a variety of collective models and their inherent properties are discussed in details, such as Vicsek model [2, 3], Cucker-Smale(CS) model [4, 5], hierarchical flocking model [6, 7, 8], metric-free model[9, 10, 11, 12], et al.

The Vicsek model, known as the simplest and the most classic model, displays a transition from random state to collective motion [2]. Subsequent numerical studies greatly helped clarify its properties on this transition [13, 14, 15, 16, 17, 18, 19, 20]. The basic principle of collective behaviour is how to form the swarm cohesion configuration for groups of individuals. Most studies either confined the whole swarm to a periodic boundary condition(PBC) [2], or introduced attraction and repulsion terms [21, 22], or some potential field [23]. Nevertheless, Pearce and Turner found that these approaches can generate swarm cohesion, but they also introduce a metric to the system [12]. These methods usually produce feature densities that are virtually constant and independent of the number of individuals in a group, which are not visible in data [24].

Most collective models like the Boid model and Vicsek model have shown that simple rules of local interaction between individuals can produce the collective behaviour of animals. Nevertheless, people know little about the nature of this interaction, so most research results rely on prior assumptions [25, 14, 26]. Ballerini and co-workers believed that the interaction does not depend on the metric distance but instead on the topological interaction mode by reconstructing the three-dimensional positions of thousands of individual birds in an airborne flock [27]. According to the experimental data, another critical topological feature of bird interactions is that each individual interacts with up to six or seven neighbours, no matter how far they are. Numerical simulations show that the topological interaction enables the swarming to achieve a better consistency state. Moreover, this interaction mode is more consistent with the actual situation of birds in nature.

Furthermore, Francesco and his co-workers found that the aggregation characteristic of self-propelled particles aligning with their topological neighbours is different from traditional metric models. In this experiment, particles only select the neighbours on the first layer of the Voronoi as the interaction objects [10]. Andrea Cavagna and his co-workers measured the correlation degree of velocity fluctuation of different birds by reconstructing each bird’s three-dimensional position and velocity in a large number of starlings, and measurement results also verified that this behaviour correlation is scale-free [9]. Rylan Wolfe et al. proposed a scale-free Vicsek model by quoting a new Enskog-type theory, and their simulation results indicated that the transition is continuous in contrast to the traditional metric Vicsek model [11]. Camperi et al. compared the stability of topological models with metric ones, and they concluded that the number of neighbours interacting with particles is constant and topological interaction mode has better anti-interference performance [28]. Nevertheless, the swarming density of most models involves topological interactions approaches zero under spatial expansion. In response to this problem, Pearce and Turner proposed a Strictly Metric-Free(SMF) swarming model that involves a metric-free motional bias on individuals topologically identified as being at the edge of swarm [12]. This model can change the density of the particle system in the unbounded region, and the motorial bias enables the system to achieve an aggregation state in the unbounded three-dimensional space. In later studies, Turner further developed this model and endowed birds at the edge of the swarm had an inward movement bias, while birds within the group must have an outward bias [29]. He extended the motional bias to act on all birds with strength prescribed by a function of the topological depth of individuals within the swarm. Experimental results show that the particles in the boundary area have a denser swarm than those in the centre area.

Couzin et al. believed that each individual in a group uses visual information as the predominant modality to interact with each other when making collective decisions [30, 31]. Most existing collective models suppose that every particle can obtain global vision information, such as the Vicsek model, Turner’s model. However, in nature, particles in a flock usually have a restricted visual field. For example, the visual field of starlings is 286°, the visual field of pigeons is 316°, while the visual field of owls is 201° [21, 32, 33]. Although it is not easy to infer how they become such a limited visual field during natural selection evolution, we can quantitatively analyze the advantage of restricted visual field compared with the global visual field.

Therefore, subsequent studies mainly focus on understanding the biological mechanism model of collective motion in nature [34, 35, 36, 37], as well as improving the convergence properties of this collective behaviour further [38, 39, 40, 41, 42, 43, 44]. Most of the existing literature mainly concentrated on reducing the convergence time of the collective evolution process of a Vicsek-like model from the point of view of interaction rules or restricted view angle. For example, Wang et al. focused on minimizing convergence time of direction consensus of a two-dimensional Vicsek model by exponential neighbour weight and restricted visual field [40]. Nguyen et al. studied the effect of visual angle on the phase transition in the two-dimensional Vicsek model, and they found that the phase transition exists when the view angle of each particle is more than *π*/2 [41]. Durve et al. believed that both the directionality and radial range of the interaction plays an essential role in determining the nature of the phase transition [42]. Li et al. optimized the three-dimensional Vicsek model with the purpose of the quickest direction consensus, and they found that the optimal view angle is concerned with the absolute velocity and the particle density [43].

Because of the above problems, this paper proposes a fully three-dimensional restricted visual field metric-free(RVFMF) model. Compared with most previous models, the RVFMF model is strictly metric-free and has a restricted visual field. Each particle chooses these particles located in the first shell of Voronoi tessellation and within its restricted visual field as its topological neighbours. This mechanism ensures that the RVFMF model is strictly scale-free. Besides, we define the individuals lying on the convex hull of the swarm as being on edge, while others are in bulk. A scale-free edge eigenvalue is introduced, which only acts on edge particles. This eigenvalue introduces a scale-free inward motorial bias for edge particles. “Inward” refers to a vector average pointing from an edge particle to its topological neighbours located on edge. Under the influence of motorial bias, edge particles can control the density of the swarm. From the predation point of view, the individual at the outermost part of the animal group is the most vulnerable to predation and will spontaneously try to move closer to the group. Meanwhile, the internal individual is relatively safe and will not move to the more dangerous cluster edge but keep the current state as much as possible.

RVFMF model can be used to simulate the evolutionary process of large-scale birds, such as starlings. We vividly restored the process of the European starlings’ flight reaching consistency through the RVFMF model and explored the influence of visual field on swarm consistency. We also try to characterize the optimal field vision of each individual under different group sizes and prove whether a global visual field is necessary for a flock.

## 2. Restricted Visual Field Metric-Free (RVFMF) Model

The RVFMF model is defined as follows. We consider *N* particles moving at the same speed in a three-dimensional borderless region. The main contribution of this model is to consider the visual angle of each particle during the evolution process of collective behaviour from initial disorder state to final order state. At the initial time *t* = 0, the position of *N* independent particles are randomly distributed in a cube with side length *L*, and the velocity direction of each particle is a random value generated from [0, 2*π*). At discrete time *t*, the position of a particle *i* is denoted as, the velocity direction is 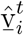, and 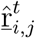 represents the unit vector pointing from particle i to particle j at time *t*.

As shown in Fig. 1, we define the visual field of each particle as the cone-like scattering column area formed by the particle. Suppose that every particle in the same group owns the same visual angle. Let *θ* ∈ (0, *π*] denote the visual angle of each particle. When *θ* = *π*, meaning that every particle has a global vision, it becomes an SMF model.

**Figure 1:**
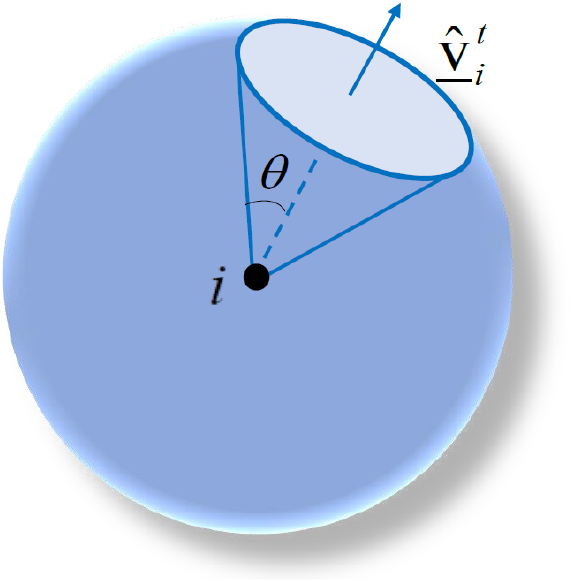
Schematic diagram of the view field *θ* of particle *i*.

According to equations (1) – (4), all particles positions and velocity directions are updated once every time step. The topological neighbours in the field of view of particle *i* belong to set *B_iv_*, which will be updated constantly according to the position of particle *i* at time *t*. All particles in the convex hull consist of set *C*. If particle *i* belongs to set *C*, then set *S_iv_* is equal to all topological neighbors of particle *i* located on the cluster convex hull, that is, *S_iv_* = B_*iv*_⋂*C*. This model has two control variables, the first parameter *ϕ_e_* is the edge strength, which is the relative weight of the inward movement deviation of edge particles, the second parameter *ϕ_n_* is the intensity of noise. See equations (1) – (4).

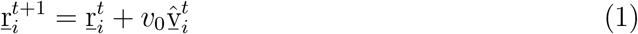

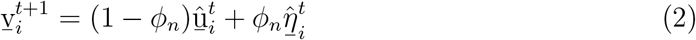

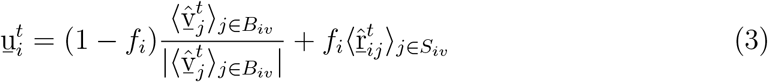

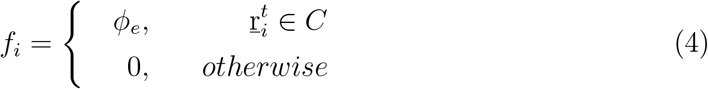

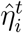 represents a random unit vector at time *t* for particle *i*, *v*_0_ represents the speed at which particles move, ^ represents the normalized unit vector, angle brackets < ··· > indicate average subsets. Because the above four equations are completely scale-free, there is no need to select specific units. The RVFMF model does not require the velocity *v*_0_ of particles to simply the cluster model. The only length unit is the distance *v*_0_ of particle movement at each moment, and the time unit is the duration of each step. In our model, the position of each particle at the next moment is equal to the current position plus the velocity of the particle multiplied by the velocity direction, that is, equation (1). Every particle suffers the same weighted noise, controlled by 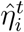 in equation (2). According to equation (3), each individual has a definite direction before introducing any noise. When particle *i* locates in bulk, *f_i_* = 0 means that it is not affected by edge strength *ϕ_e_*, and its direction are equal to the average direction of its topological neighbours. When particle *i* is on edge, *f_i_* = *ϕ_e_* means that particle is direction is equal to the linear superposition of the direction average of its topological neighbours multiplied (1 − *ϕ_e_*) and the scale-free motorial bias multiplied by *ϕ_e_*.

The model involved here is in three-dimensional space. For better explanation and clearer exhibition, Fig. 2 gives a schematic diagram of the topological constructions in equation (2) – (3). The purple dots denote the particle 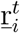 located on the convex hull, while the red dots denote the particle 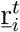 located in bulk. In order to build Voronoi tessellation, Delaunay triangulation must come first, which is represented by a red line. The meaning of point set *B_iv_* is the topological neighbours within the visual field of the target particle, which is composed of all particles linked by a red line and purple line from the target particle. The purple line connects the particles on the convex hull, and in the point set *S_iv_* = *B_iv_*⋂*C*, pink arrows indicate the scale-free inward motorial bias of these edge particles. Fig. 3 shows more details on the composition of the metric-free surface term appearing in equation (2) – (3) for edge particle *i*. Metric-free surface term of edge particles are equal to the average value of unit vectors 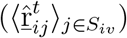 of topological neighbors which are also located on convex hull, that is, the average of the unit vectors 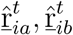 of adjacent particles *a* and *b*. This value has a magnitude in [0,1] when the angle between particles *i* and *j* is small. The longer the arrow is, the larger the value is. Therefore, the larger the surface term of the outermost particles in the convex hull is, the stronger the trend of re-entering the swarm is. Moreover, to show the convex hull structure and topological neighbour structure of population more clearly, the structure of the swarm convex hull and the composition of topological neighbours in three-dimensional space are shown in Fig. 4. Therein, neighbours located within the field of view of the selected particle are represented by different colours, highlighting the restricted visual field of our model.

**Figure 2:**
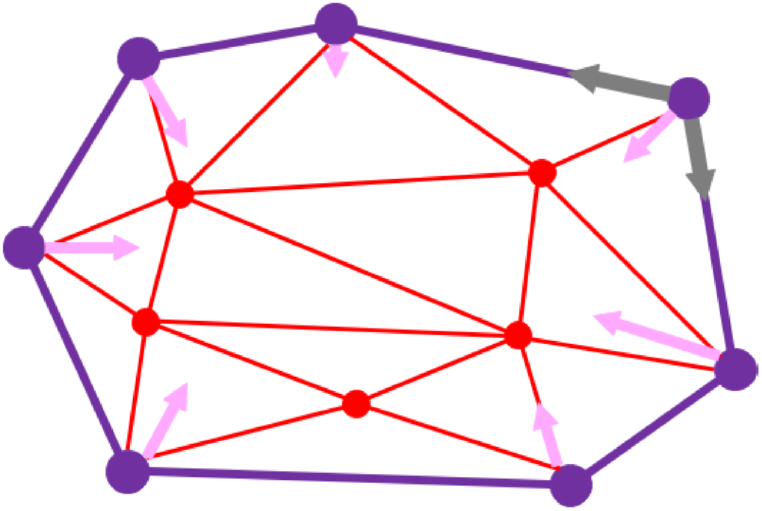
A schematic diagram on the topological constructions of particles.

**Figure 3:**
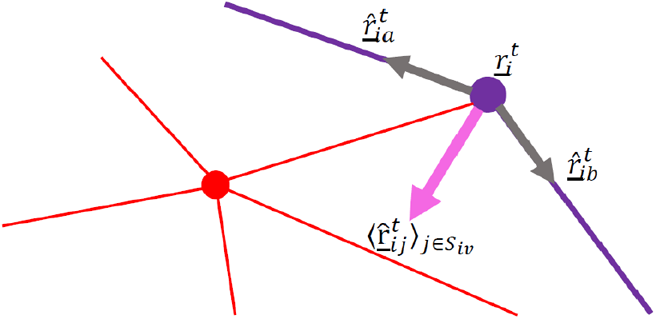
Details on the topological structure of edge particles on convex hull.

**Figure 4:**
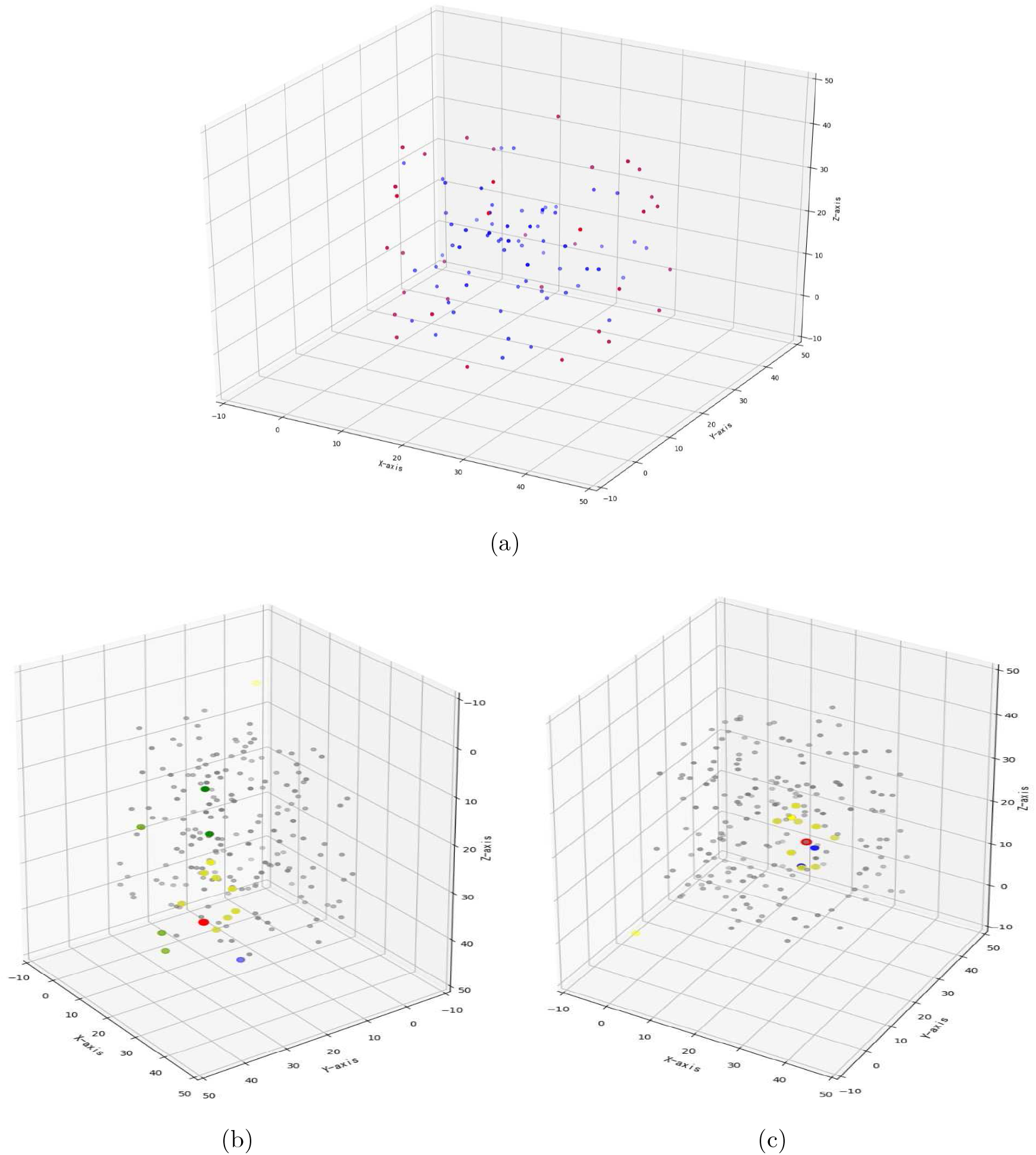
Convex hull and topological neighbor structure. (a) Red particles are located on the convex hull of the swarm, while blue ones are located in the bulk. (c) The red particle is randomly selected, blue particles are its ordinary neighbors, and topological neighbors inside the field of view is yellow. (b) If the selected particle is located on the convex hull, its topological neighbors on the convex hull is green, other neighbor colors are the same as in (c).

In order to describe the consistency degree of swarm, this paper introduces the concept of order degree

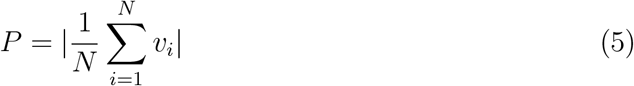

Order degree *P* represents the consistency degree of all individuals in the swarm, equal to the average centre of the mass speed of the swarm. The value of *P* falls on [0, 1]. The larger the *P* are, the better the consistency of these particles is. When *P* = 1, the movement direction of all particles is consistent, similarly, at the initial moment of random distribution of particles, *P* = 0.

## 3. Simulations Analysis and Results

European staring is a ground-eating bird of the order Passeriformes that moves in the daytime [45]. Inspired by their flocking behaviour, a three-dimensional collective model called RVFMF is developed here to investigate the influence of the field vision on the whole systems stability when approaching a flock. In order to exhibit the evolutionary process of the system from disorder state to order state, a human-computer interaction interface shows the simulation process of the RVFMF model under different parameters, such as population size *N*, noise intensity *ϕ_n_*, and edge intensity *ϕ_e_*. Fig. 5 provides an example to present the main interface contents of the simulation platform, from which we can observe the changing process of a cluster from an initial state (as shown in Fig. 5(a)) to a stable state (as shown in Fig. 5(b)). The colour of each particle denotes its velocity direction. The same colour means the same heading direction. Fig. 5(c) shows the curve of the global order *P* of the system over time.

**Figure 5:**
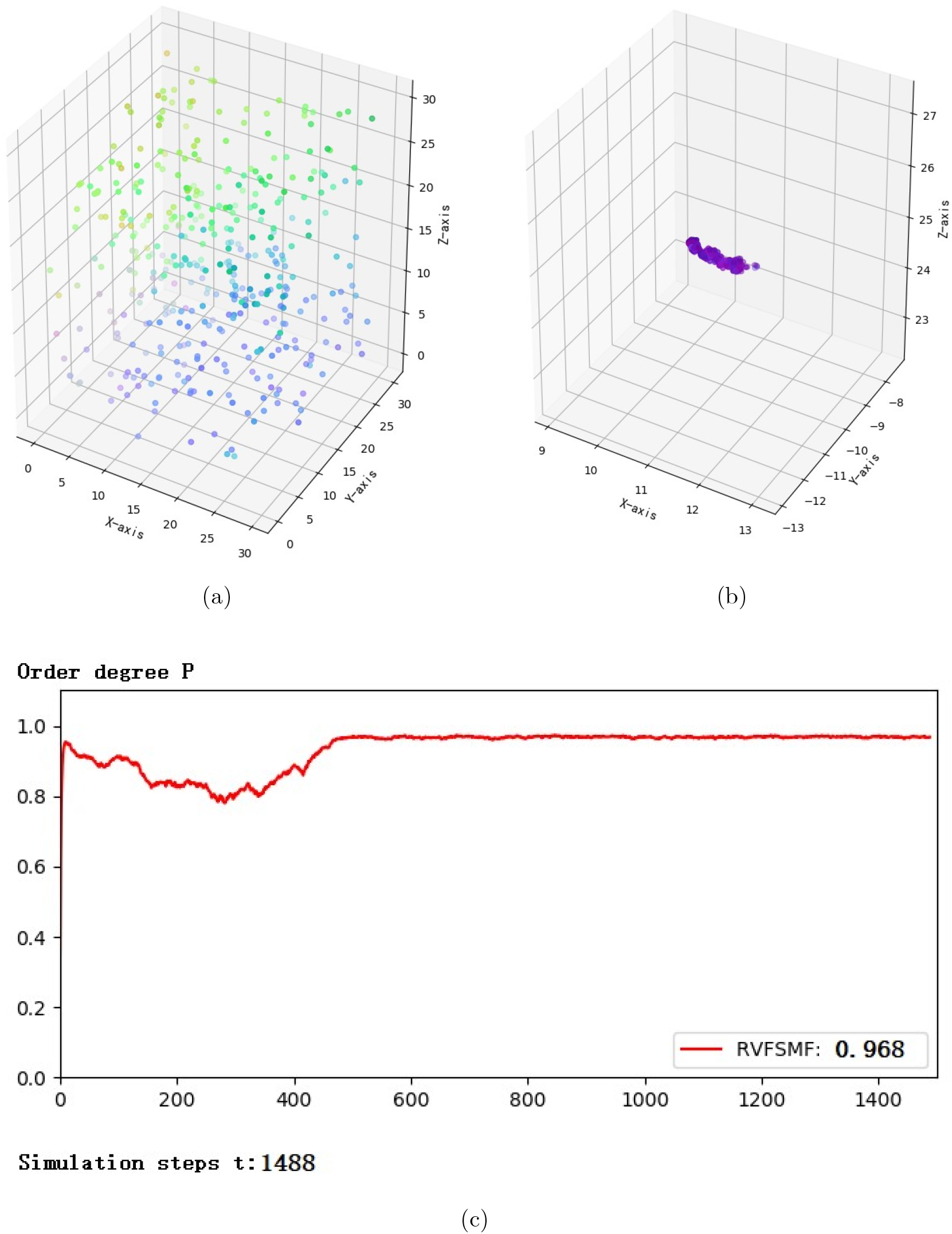
Swarm from disordered state to ordered state, in which we set the system as a global vision, i.e. *θ_i_* = *π*, *i* = 1, ···, *N*. Other parameters are population size *N* = 400, noise *ϕ_n_* = 0.2, *ϕ_e_* = 0.5. (a) Initial state of random distribution. (b) Stable state of overall speed consistency. (c) Global order *P* of swarm tends to be stable from 0 to 0.968. (The color of the particles in the picture represents the speed direction of the particles, the closer the color is, the higher the global order *P* of the swarm is.)

### 3.1. Two control parameters

Our purpose is to study the influence of the visual field of each particle on the swarming system’s stability. In order to explore its influence, we need to control other variables to remain unchanged. Our model has two control parameters, the weight of inward motion bias of particles on the edge *ϕ_e_* and the size of noise *ϕ_n_*.

First, we change the two control parameters separately and keep the other variables unchanged. The goal is to observe the influence of the two parameters in the global order *P*. Suppose each particle owns a global vision, observe the changes of the global order of swarm with different group sizes affected by noise strength *ϕ_n_*, and Fig. 6 shows the simulation results. The results show that swarm can achieve a higher degree of consistency only when the noise interference is minor, and the higher the noise, the lower the consistency of the swarm, and the change of population size will not change this trend. These results are the same as those in the reference [12], which also verifies the correctness of our model; that is, when each particle owns a global view, the RVFMF model becomes an SMF model.

**Figure 6:**
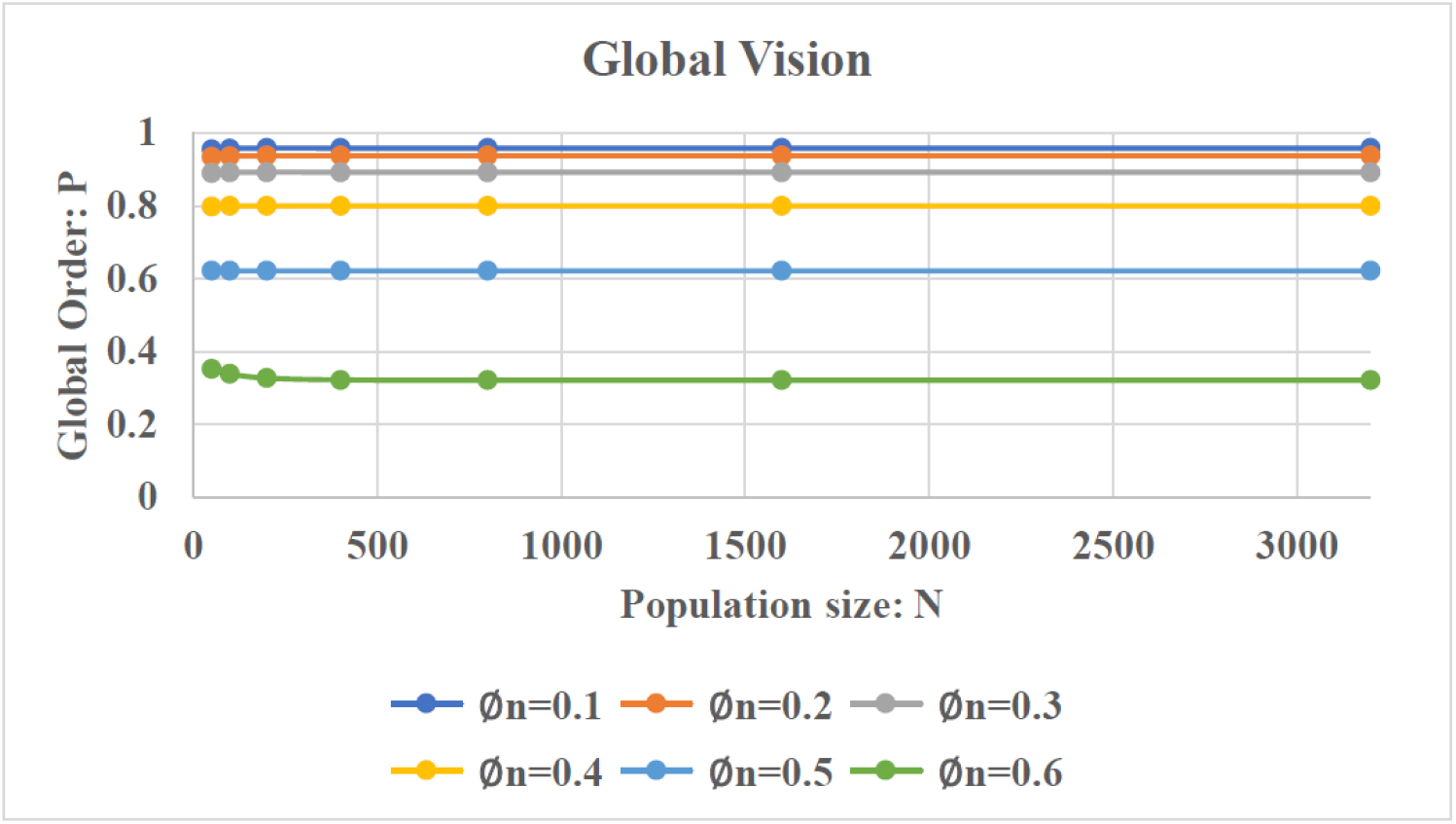
Under the global view, the swarm is disturbed by different degrees of noise, legend of trend of global order changing with population size. We keep *ϕ_e_* = 0.5.(Providing the same weight of synergy and inward bias for individuals on the boundary.)

Next, under the global vision, observe the change of the global order of swarm with different group sizes affected by edge strength *ϕ_e_*, and Fig. 7 shows the simulation results. From the results, we observed that when the population size did not reach 1000, the change of *ϕ_e_* had an evident impact on *P*, as the population size continued to increase, the change of *ϕ_e_* value had little effect on *P*. When the population size is small, the impact is that the higher the *ϕ_e_*, the lower the order degree that the swarm can achieve. According to equation (3), the trend of co-aligning with neighbours and re-entering the swarm being weighted by factors (1 – *ϕ_e_*) and *ϕ_e_* respectively. The larger the *ϕ_e_* value is, the smaller the weight of cooperation is, so the smaller global order *P* is. Furthermore, under the same population size and the same noise strength, the stronger the edge is, the worse the systems stability. When the edge strength is no more than 0.5, the systems stability is almost the same with different population size. On the contrary, when the edge strength is more significant than 0.5, under the same edge strength, the larger the group size is, the better the system’s stability is. Therefore, we will mainly discuss how to optimize the visual field angle according to the systems stability when the edge strength is more significant than 0.5.

**Figure 7:**
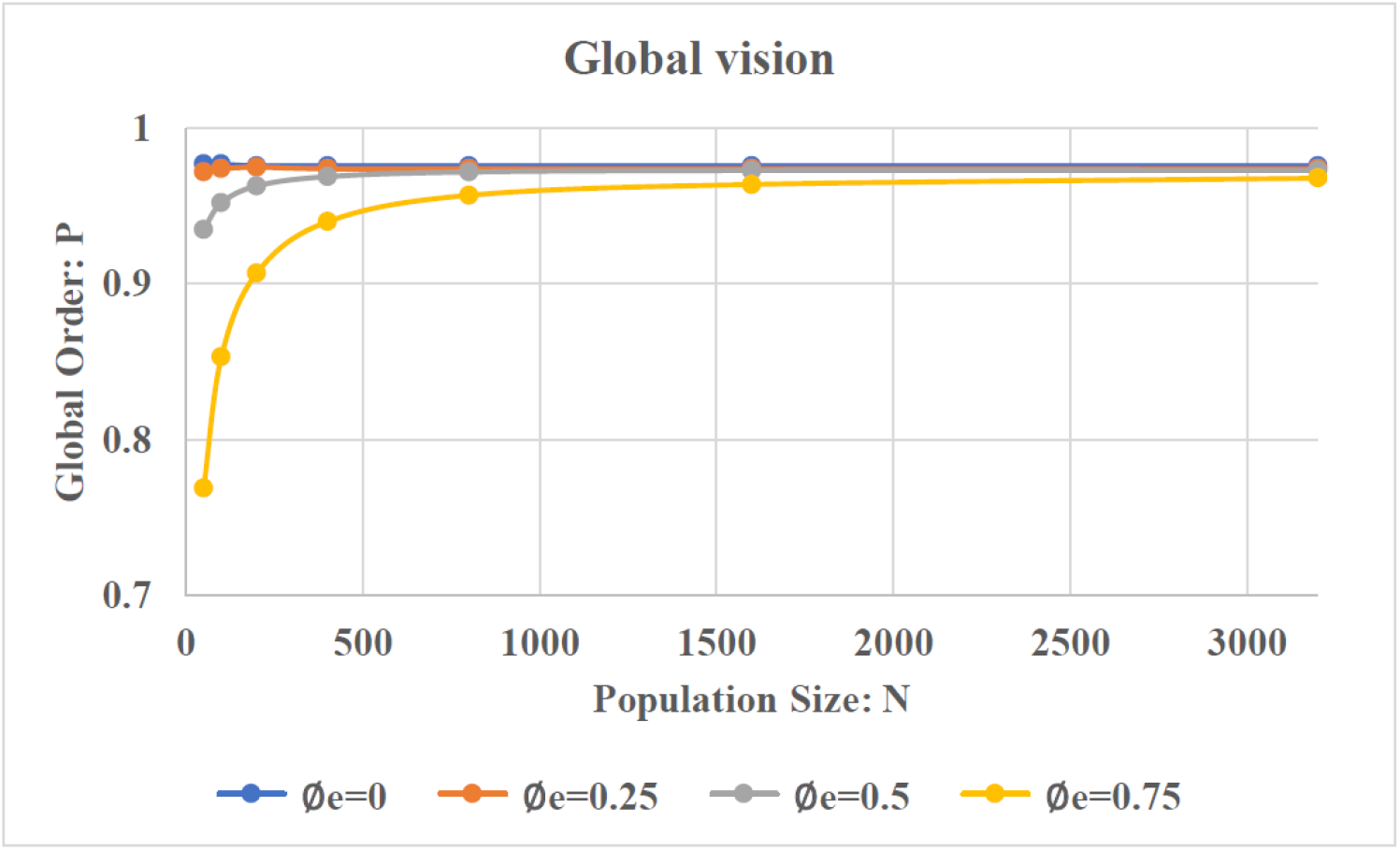
Under the global view, the swarm is disturbed by different degrees of *ϕ_e_*, legend of trend of global order changing with population size. We keep *ϕ_n_* = 0.2.

### 3.2. Influence of visual field angle

In general, people think that the global angle can provide individuals with the largest field of vision and sufficient information, so the system can achieve the highest degree of cooperation when *θ* = *π*. At last, however, the related research on European starlings in biology have pointed out that the eyeball of European starlings is in a specific limited field of vision rather than a global visual field during flight, and when the starlings are at rest, the visual field is between 130° and 160° in the vertical direction[45]. Why they act in this way may be determined by complex biological mechanisms. In this paper, we would like to investigate whether the restricted visual field angle will benefit the stability of a flock or not according to quantitative simulation analysis.

Considering that the starling’s eyeball can rotate during flight, which will lead to a 20° margin in the vertical direction, we assume that the starling’s visual field angle ranges from 110° – 180°. In order to observe the influence of visual field angle on the order degree of a swarm, we control other parameters to be constant and set the visual field angle from 110° to 180° at intervals of 10°.

Based on the previous analysis, we would like to discuss how to choose the edge strength *ϕ_e_* and noise strength *ϕ_n_* firstly. Let the population size *N* = 500 and the noise strength *ϕ_n_* = 0.3. Fig. 8 shows the curve of global order *P* under different edge strength *ϕ_e_* and different visual field angle. As shown in Fig. 8, when *ϕ_e_* = 0.25, 0.35, 0.4, that is, when *ϕ_e_* ≤ 0.5, the global order *P* increases with the increasing field of view angle *θ*, which shows that when the proportion of co-alignment is high, the influence of the field of view angle is not apparent to the starling swarm. On the contrary, when *ϕ_e_* > 0.5, we can see that with the increase of visual field angle, *P* owns its maximum value.

**Figure 8:**
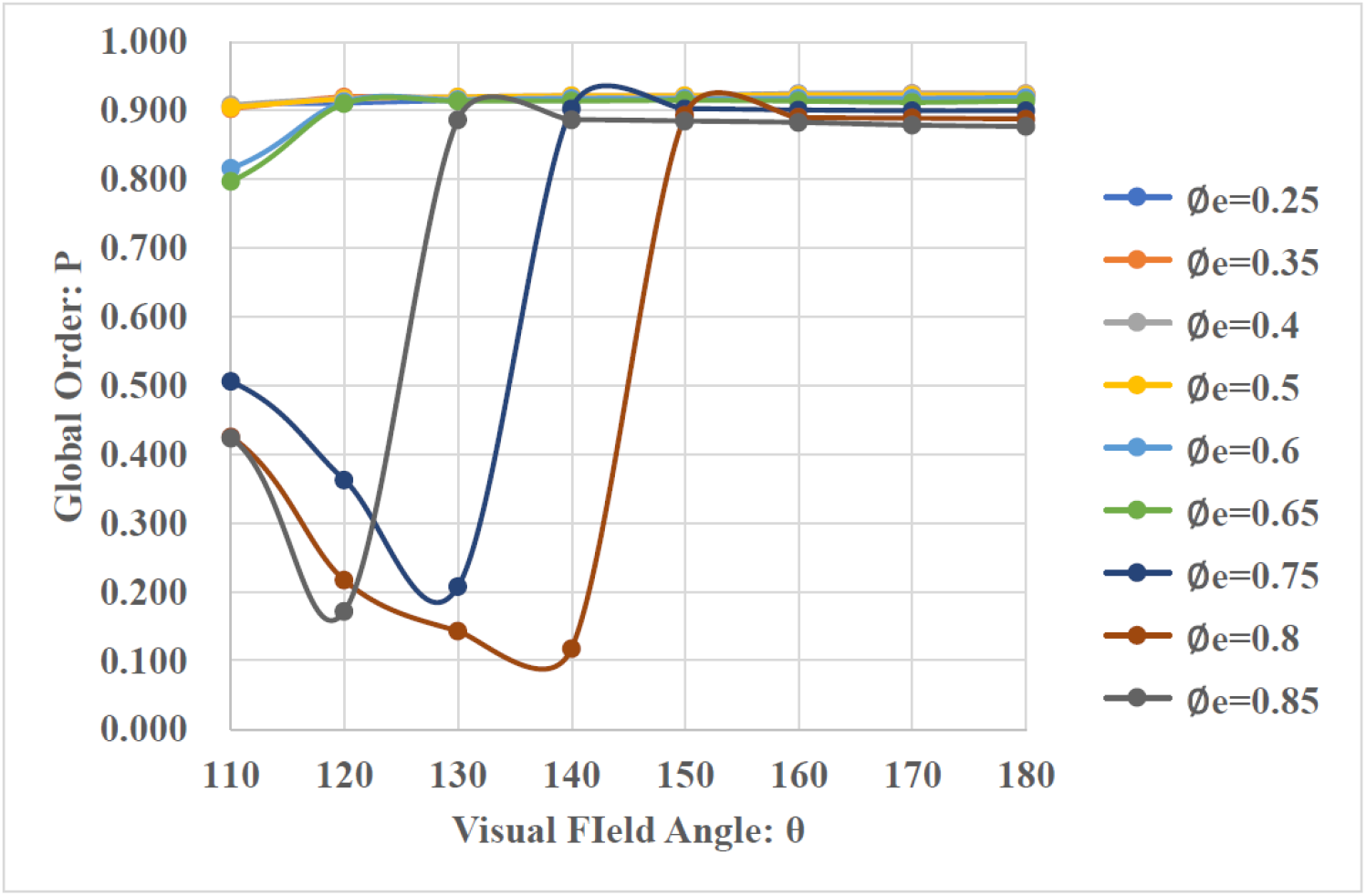
*ϕ_e_* takes different values, and global order *P* changes with visual field angle. Other parameters are *N* = 500, *ϕ_n_* = 0.3.

Without loss of generality, we finally choose *ϕ_e_* = 0.75, *ϕ_n_* = 0.3 as experimental parameter. Population size *N* takes different values, the change of global order *P* of the system with visual field angle is shown in Fig. 9. As shown in Fig. 9, under different population sizes, we can observe a visual angle in the starling swarm system, making the swarm finally achieve the highest global order. We define this angle as the best view angle and mark it with *θ_opt_*. This angle is less than 180°, which is not a global view.

**Figure 9:**
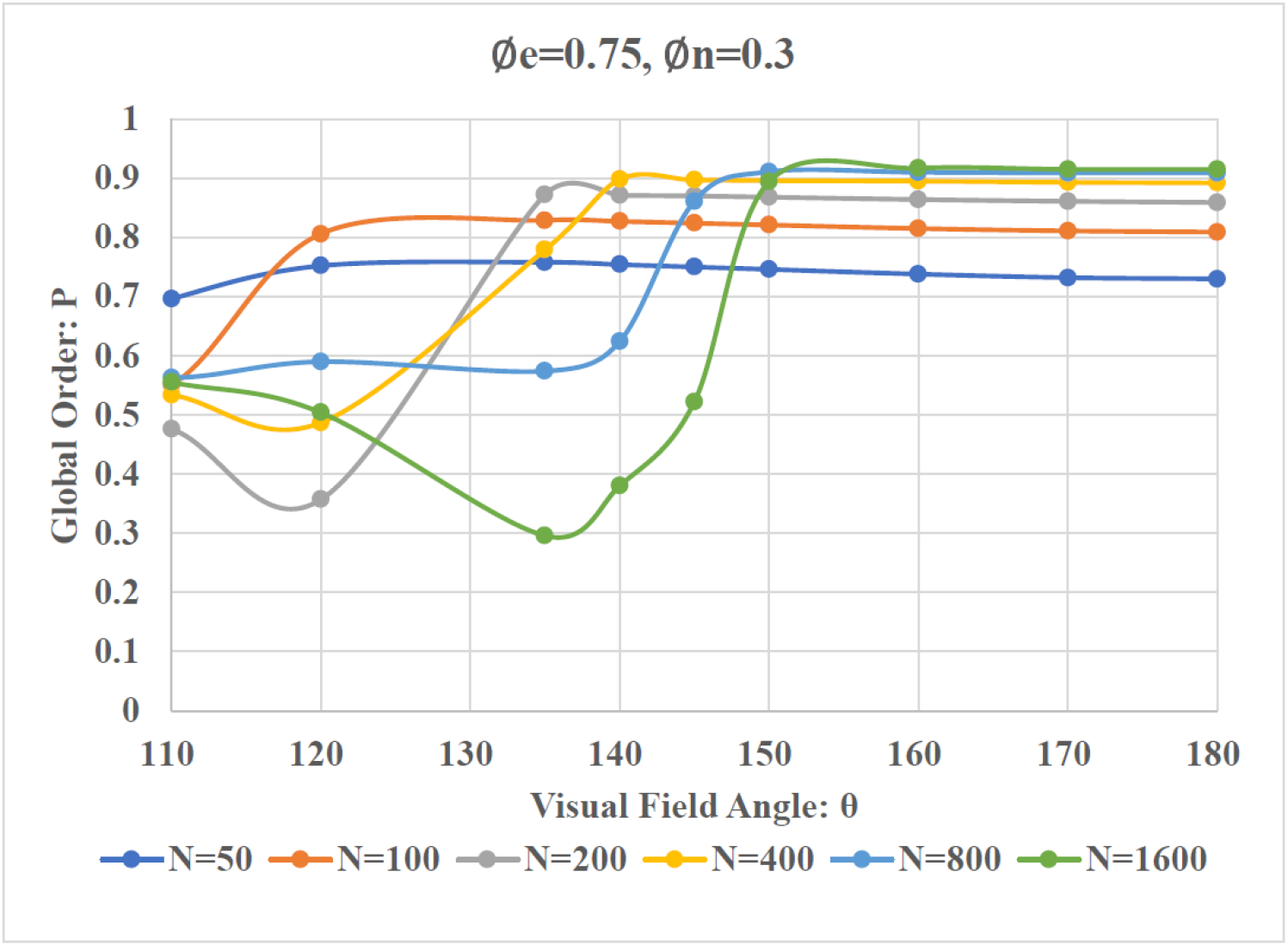
*N* takes different values, and global order *P* changes with visual field angle(When the value of *ϕ_e_* is large, it is difficult for the system to reach a stable state when the number of iteration steps is small. Therefore, after 10000 pre-balancing steps, the average value of the global order *P* of 1000 steps is selected as experimental data, and the experiment is repeated 100 times to take the average value.)

In order to further explore the relationship between the best view angle *θ_opt_* and the population size of the system *N*, plenty of simulation experiments are carried out under the condition that *ϕ_e_* = 0.75, *ϕ_n_* = 0.3, and keep the visual field angle from 110°to 180°at intervals of 5°. Fig. 10 shows the best view angle of the swarm under population size. The above simulation results conclude that the best view angle of the starling swarms increases almost linearly with the increase of population size, and *θ_opt_* tends to be stable when the population size is more significant than 1000. It seems that the size scale 1000 is a critical inflection point, which is worthy of being developed further by scientists.

**Figure 10:**
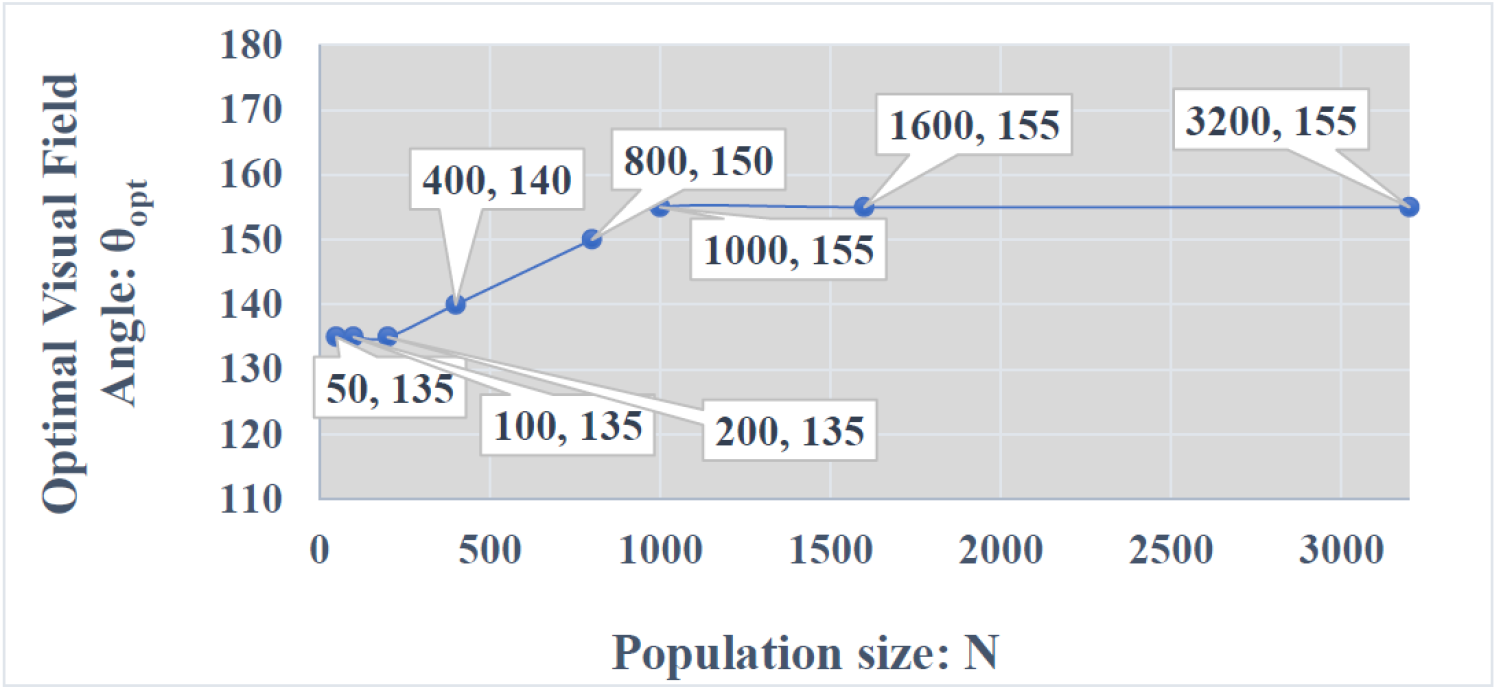
The best view angle changes with the population size.(The system conducted 100 experiments under different field of view of each population size, and global order *P* of each experiment corresponds to an average over 1000 simulation time steps following a 10000 time step pre-equilibration period.)

### 3.3. Discussion and conclusion

In Fig. 6, we observe that swarms of different group size can achieve the same degree of order parameter under different degrees of noise interference. In nature, there is often such a phenomenon that a large European starling swarm suddenly divides into two swarms during flight and keeps the same order to continue flying. The results in Fig. 6 explain this phenomenon. Under the same degree of noise interference, individuals in the swarm can keep the aggregation state unchanged without being affected by population size.

Fig. 7 shows that the order degree of the system is concerned with the inward bias degree of individuals who are on the edge when the population size is small enough (such as no more than 1000). It is also easy to understand that with the increase of inward bias weight, cooperation is lower, and it is more difficult for swarms to achieve overall speed consistency. At the same time, we find that with the increase of the population size, the influence of *ϕ_e_* becomes smaller and smaller. That is to say, the larger the population size is, the more stable it is, and the less it depends on the edge strength. That is why European starlings tend to form a larger whole for activities, which is very safe and stable.

Existing literature has referred that the eyeball of European starlings is in a specific limited field of vision rather than a global visual field during flight [45]. As shown in Fig. 8, the experimental results when *ϕ_e_* > 0.5 are quite consistent with this conclusion. To some extent, experimental data in Fig. 8 also validate that in the actual starlings, the individuals on edge are more inclined to return to the swarm instead of cooperating with the neighbours. This tendency effectively guarantees that these individuals do not break away from the swarm.

According to the experimental results in Fig. 9, we concluded that the starlings have an optimal field of view. The experimental results in Fig. 10 show that the best view angle of the swarm will increase with the expansion of the population size, and the best view angle tends to converge and stabilize around 155° as the population size reaches around 1000. When the swarm does not reach a specific scale(such as 1000), individuals need to appropriately increase and adjust their visual field to maintain a more efficient interaction with the expansion of the population size.

Suppose each particle adopts the optimal vision as shown in Fig. 10, observe the changes of the global order of swarm with different group sizes affected by noise strength *ϕ_n_*, and the results are shown in Fig. 11. Furthermore, we compared the global order *P* under global vision with that under optimal vision. Fig. 12 shows the experimental results. It is clear to see that when the noise strength *ϕ_n_* is smaller than 0.5, then the system with optimal vision approaches to a higher consistency finally. Nevertheless, when the noise strength *ϕ_n_* is more significant than 0.5, then global vision works better. Consider that noise strength is usually no more than 0.5. Therefore, during the swarm flight, individuals tend to evolve into such an optimal visual field to maintain overall stability, involving a more complicated biological mechanism. Moreover, optimal vision seems to be a better solution in directional flocking or swarming behaviours.

**Figure 11:**
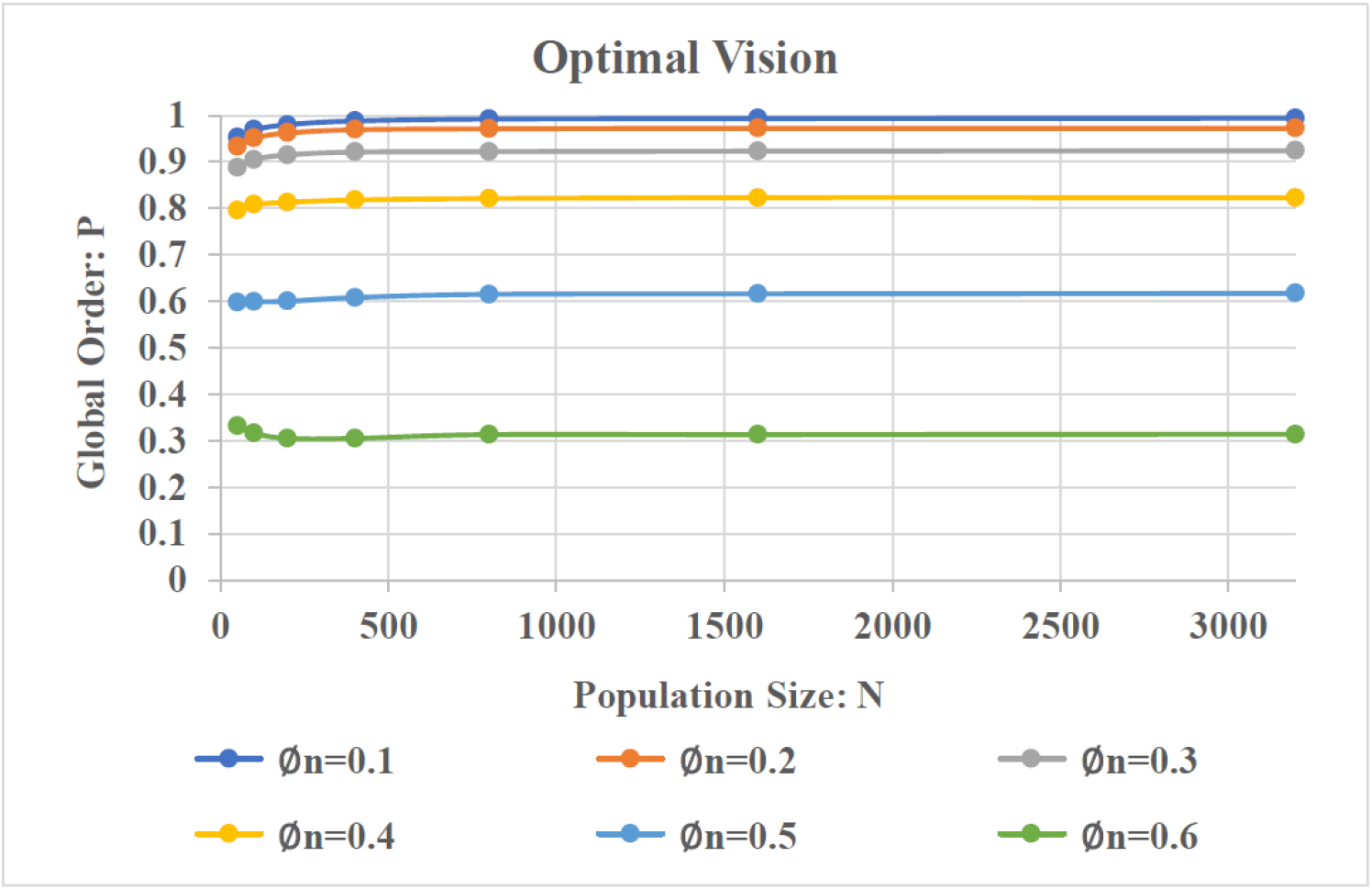
Consider each particle owns the optimal vision angle. The relationship between global order *P*, noise strength *ϕ_n_*, and population size *N*. Here, *ϕ_e_* = 0.75.

**Figure 12:**
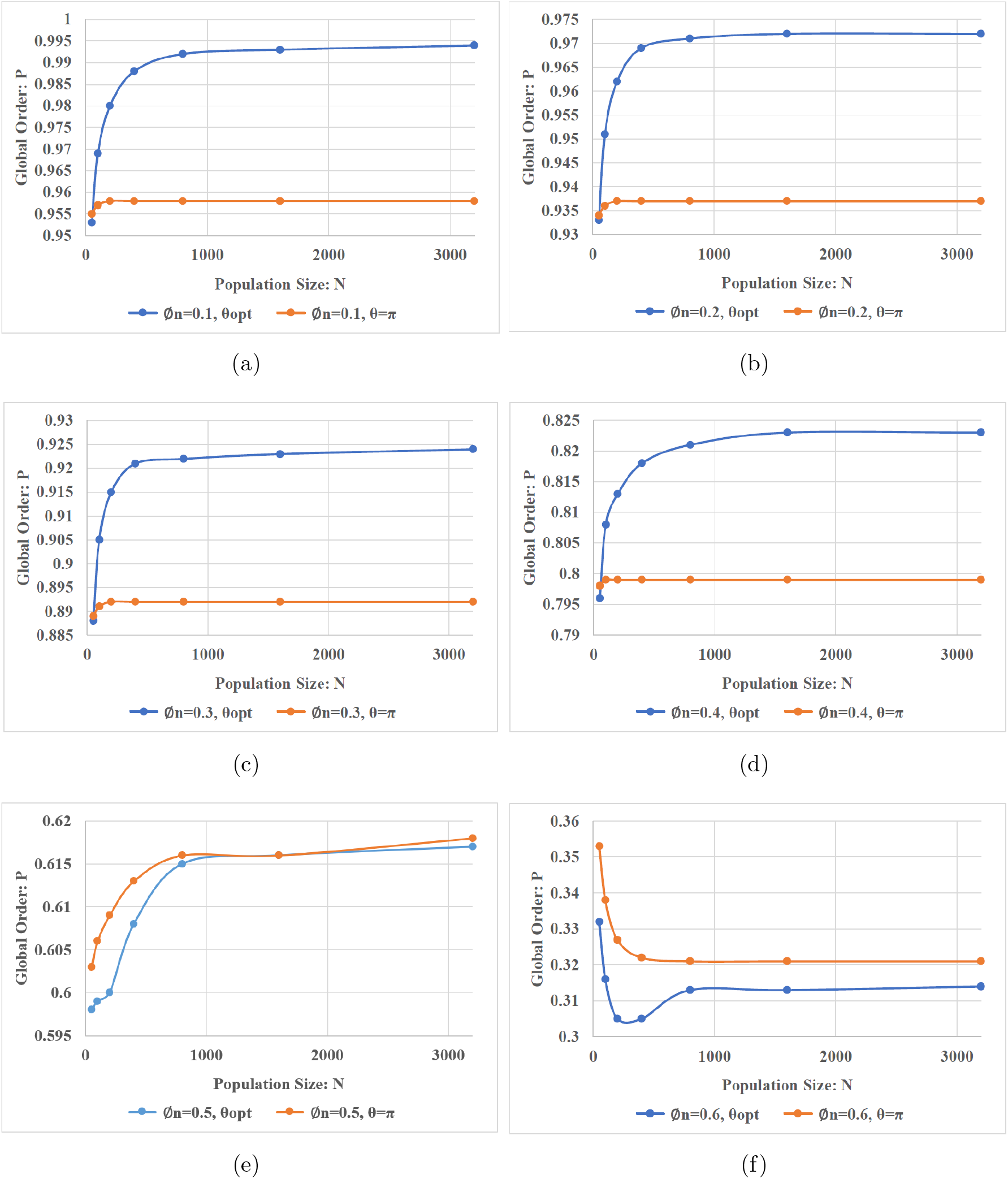
Quantitative comparison on the optimal vision and global vision according to the stability of the swarm system under different noise strength.

## Acknowledgements

This work was supported by National Natural Science Foundation of China, and Fundamental Research Funds for the Central Universities.

## Author contributions

**Conceptualization**: Yongnan Jia

**Data curation**: Lingwei Zhang

**Formal analysis**: Yongnan Jia, Lingwei Zhang, Tianzhao Lu

**Funding acquisition**: Qing Li

**Investigation**: Yongnan Jia

**Methodology**: Yongnan Jia

**Project administration**: Qing Li

**Resources**: Yongnan Jia

**Software**: Tianzhao Lu, Lingwei Zhang

**Supervision**: Qing Li

**Validation**: Qing Li

**Visualization**: Tianzhao Lu, Lingwei Zhang

**Writing-original draft**: Lingwei Zhang

**Writing-review and editing**: Yongnan Jia

## Data availability statement

All data that support the findings of this study are included within the article (and any supplementary files).

